# Iron-sulfur cluster proteins present the weak spot in plasma-treated *Escherichia coli*

**DOI:** 10.1101/2025.01.08.631878

**Authors:** Marco Krewing, Kim Marie Weisgerber, Tim Dirks, Ivan Bobkov, Britta Schubert, Julia Elisabeth Bandow

## Abstract

Non-thermal atmospheric pressure plasmas have an antiseptic activity beneficial in different medical applications. In a genome-wide screening, hydrogen peroxide and superoxide were identified as key species contributing to the antibacterial effects of plasma while [FeS] cluster proteins emerged as potential cellular targets. We investigated the impact of plasma treatment on [FeS] cluster homeostasis in *Escherichia coli* treated for 1 min with the effluent of a microscale atmospheric pressure plasma jet (µAPPJ). Mutants defective in [FeS] cluster synthesis and maintenance lacking the SufBC_2_D scaffold protein complex or desulfurase IscS were hypersensitive to plasma treatment. Monitoring the activity of [FeS] cluster proteins of the tricarboxylic acid cycle (aconitase, fumarase, succinate dehydrogenase) and malate dehydrogenase (no [FeS] clusters), we identified cysteine, iron, superoxide dismutase, and catalase as determinants of plasma sensitivity. Survival rates, enzyme activity, and restoration of enzyme activity after plasma treatment were superior in mutants with elevated cysteine levels and in the wildtype under iron replete conditions. Mutants with elevated hydrogen peroxide and superoxide detoxification capacity over-expressing *sodA* and *katE* showed full protection from plasma-induced enzyme inactivation and survival rates increased from 34% (controls) to 87%. Our study indicates that metabolic and genetic adaptation of bacteria may result in plasma tolerance and resistance, respectively.

Graphical abstract

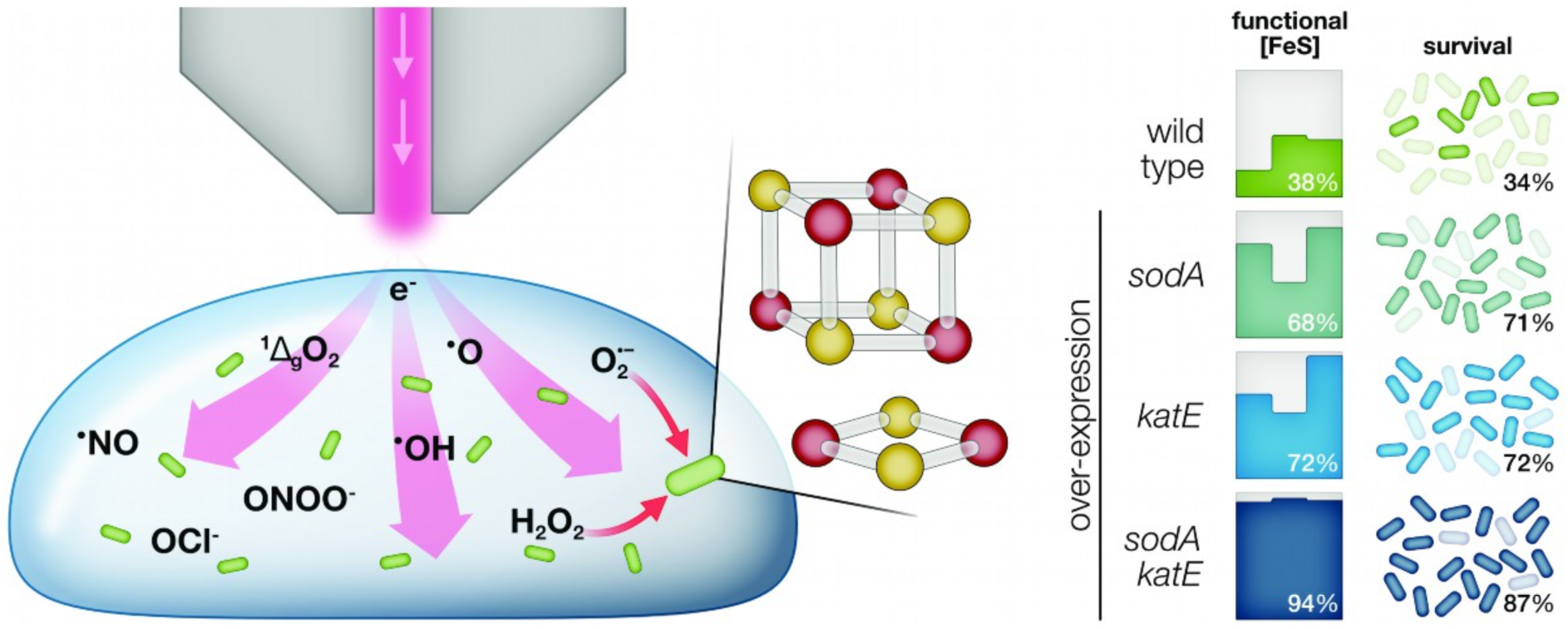

**Highlights:**

- Enzymes with [FeS] clusters are rapidly inactivated by plasma
- Clusters damaged by plasma are repaired *in vivo* using iron and cysteine
- Over-expression of *sodA* and *katE* completely prevents disruption of [FeS] clusters by plasma
- Plasma resistance is increased threefold by SodA and KatE over-production
- Pre-adaptation of *E. coli* to O_2_^-^ increases plasma tolerance

## 1. Introduction

Cold atmospheric-pressure plasmas are used in biology and medicine for sterilization, disinfection, and decontamination utilizing their antimicrobial properties ^1–5^. Plasmas are generated by means of electric fields ionizing defined gas mixtures or ambient air. In the gas phase ^6^ and in contact with liquids, various reactive oxygen and nitrogen species are generated, *e.g.* superoxide (O_2_^-^), hydrogen peroxide (H_2_O_2_), hydroxyl radicals (^•^OH), nitric oxide (^•^NO), peroxynitrite (ONOO^-^), and hypochlorite (OCl^-^) ^7–11^. Those species target biomolecules in cells causing a rapid inactivation of bacteria ^7, 12–14^.

Defects in the cell envelope ^8, 12, 15^, the DNA ^14, 16^, or proteins in key metabolic pathways ^17^ have been observed. Given the complexity of the cocktail of plasma-generated species and the additional contributions of (V)UV radiation and electric fields, the mode of action is still under investigation. It is unclear if survival hinges on a particular weak link in the chain, or if the accumulation of damage to different cellular components causes cell death. It has been shown, *e.g.*, that proteins suffer from plasma-induced modifications on amino acid side chains and from degradation into peptides resulting in rapid protein inactivation ^13, 18–20^. The sulfur-containing amino acids methionine and cysteine have been identified as most susceptible to oxidation and nitrosylation ^19, 21^.

In a genome-wide screening for plasma-sensitive mutants of *Escherichia coli*, genes and their corresponding proteins were identified, which are crucial for surviving short exposures to the effluent of a helium/oxygen plasma. Approximately 20% of the proteins are involved in iron and cysteine metabolism or the biosynthesis and maintenance of iron-sulfur clusters ([FeS] clusters) ^22^. Combined transcriptomic and proteomic studies of *Bacillus subtilis* characterized the effects of exposure to an argon plasma. The identified differentially expressed genes and differentially abundant proteins also implied iron and [FeS] clusters as determinants of plasma resistance ^23^.

[FeS] clusters are crystal-like structures of iron and sulfide arranged in rhombs or cubes. In *E. coli*, [FeS] clusters are synthesized either by the *isc* or the *suf* pathway. The *isc* operon is expressed constitutively, whereas the *suf* operon is induced upon oxidative stress and iron limitation ^24^. [FeS] clusters are the most abundant prosthetic group in proteins and participate in a wide range of processes such as respiration or DNA repair ^25, 26^. Depending on the reaction type, [FeS] clusters can either act as Lewis acid and undergo transient covalent bond formation with the substrate or they participate in electron-flow reactions as redox-cycling cofactors. When acting as Lewis acid, like in aconitase or fumarase of the citric acid cycle, [FeS] clusters are exposed to the solvent, while redox-cycling clusters, like in succinate dehydrogenase, are coordinated inside the protein and shielded from the solvent ^27, 28^. Despite their widespread occurrence and importance to many pathways, [FeS] clusters are easily damaged by molecular oxygen and reactive oxygen or nitrogen species. Superoxide and hydrogen peroxide rapidly oxidize [FeS] clusters causing disassembly and release of iron ions ^29^.

Enzymes harboring [FeS] clusters are involved in many central metabolic pathways but highly prone to oxidative inactivation. Based on the large number of [FeS] cluster-related genes identified in the two global studies on the effects of plasma exposure on bacteria mentioned above ^22, 23^, we hypothesized that [FeS] clusters might present the weak link in bacterial cells exposed to plasma. On that account, we investigated to what extent [FeS] cluster-harboring enzymes are affected by plasma. We then tested in how far their protection improved survival under plasma exposure, investigating the physiological effects of iron supplementation on plasma tolerance as well as consequences of plasmid-based over-expression of protective enzymes on plasma resistance.

## 2. Material and methods

### Plasma source

Plasma was generated using a micro-scale atmospheric-pressure plasma jet (µAPPJ) ^30^. The feed gas was a mixture of helium (1.4 slm, 5.0 purity) and oxygen (0.6%, 4.8 purity). The plasma was ignited at 13.56 MHz and 230 V_RMS_. For plasma treatment, 20 µl of aqueous samples were applied onto a filter paper (0.9 cm in diameter, Whatman, 3MM) and placed at a 4 mm distance to the nozzle of the jet.

### Strains, media, and chemicals

*E. coli* BW25113 was used as wild type. All corresponding deletion mutants were derived from the KEIO collection of single-gene knockout mutants ^31^. Plasmids for over-expression originated from the ASKA collection ^32^. The double deletion mutant Δ*sodA*Δ*sodB* (Δ*sodAB*) was generated by removing the kanamycin resistance cassette replacing the *sodA* gene in the Δ*sodA* strain and then inserting a kanamycin resistance cassette in the *sodB* locus using P1 phage transduction and standard protocols ^33^.

Following the same procedure, the deletion strains Δ*sodA*Δ*sodB*Δ*sodC* (ΔSOD), Δ*katE*Δ*katG* (ΔKAT), and Δ*katE*Δ*katG*Δ*ahpC* (ΔHPX) were generated. To construct the plasmid pCA24N::*sodA*::*katE* for *sodA* and *katE* over-expression, the gene *katE* was amplified by PCR using the primers 5’-aaaagtcgaccaattgtgagcggataac-3’ and 5’-aaaagtcgacaggagtccaagctcag-3’ and the plasmid pCA24N::*katE* as template. After digestion with *Sal*I, the fragment was ligated into pCA24N::*sodA*. All cells were cultivated in LB medium or M9 minimal medium at 37°C under aerobic conditions and inoculated from overnight pre-cultures. Kanamycin (50 µg ml^-1^) or chloramphenicol (50 µg ml^-1^) were used when necessary.

Gene expression from plasmids was induced by isopropyl thiogalactoside (IPTG), which was added directly at inoculation of the main culture. Protein expression was measured using SDS-PAGE and Western blotting following to standard protocols. To this end, 1 ml of bacterial culture was harvested at an OD_600_ = 0.6, resuspended in 75 µl denaturing loading buffer, and subsequently heated to 95°C for 10 min. For SDS-PAGE, 5 µl of each sample were applied. After Western transfer, His_6_-tagged proteins were detected with a DyLight649-conjugated anti-His_6_ antibody (antibodies-online, Aachen, Germany) using the ChemiDoc MP Imaging system (Bio-Rad, Germany). Band intensities were quantified with the Image Lab software (Bio-Rad, Germany)

### Determination of plasma sensitivity

Plasma sensitivity was determined as described previously ^22^. In short, bacterial cultures in logarithmic growth phase (OD_600_ = 0.3-0.6) were first diluted to an OD_600_ of 0.3 and then 1:5,000 in LB medium to give app. 20,000 CFU ml^-1^. 20 µl (∼400 CFU) were applied onto a filter paper and exposed to the effluent of the µAPPJ for 30 s. Plasma was ignited or not, resulting in plasma-treated samples and gas controls, respectively. Directly after treatment, the filter paper was immersed in 1 ml NaCl (0.9% (w/v)) and mixed vigorously. The paper was removed, and the cell suspension plated onto LB agar for colony counting next day. CFU counts of the plasma-treated samples were set in relation to the corresponding gas controls yielding the survival rates. The plasma treatment time was set to 30 s, resulting in survival rates for the wild type without any empty vector between 0.4 and 0.5.

### Enzyme assays

Activity assays for three [FeS] cluster-containing enzymes (aconitase, fumarase, and succinate dehydrogenase) were performed to provide a readout on *in vivo* cluster integrity and functionality. The activity of malate dehydrogenase, which does not contain [FeS] clusters, served as control. Enzyme activities were determined in *E. coli* cell extracts obtained by sonication. While for the succinate dehydrogenase activity assay crude extracts were used, for aconitase, fumarase, and malate dehydrogenase, extracts were cleared by centrifugation. Protein concentrations were determined by Bradford assay ^34^ and defined amounts of protein were used in the activity assays.

Aconitase (ACN) activity was determined by following the conversion of isocitrate to *cis*-aconitate at 240 nm in Tris buffer (50 mM, pH 8) containing 10 mM MgCl_2_ ^35^. 120 µg ml^-1^ protein from clear *E. coli* lysate were applied. After 2 min pre-incubation, the reaction was started by addition of isocitrate (10 mM).

Fumarase (FUM) activity was determined by monitoring the conversion of malate to fumarate at 250 nm in Tris buffer (50 mM, pH 7.4) containing 100 mM NaCl ^36^. 120 µg ml^-1^ protein from clear *E. coli* lysates were applied and the reaction started by addition of malate (50 mM) after 2 min pre-incubation. Succinate dehydrogenase (SDH) activity was measured in crude cell lysate, using 120 µg ml^-1^ protein in Tris buffer (50 mM, pH 7.4) containing 0.1% (v/v) triton X-100. Dichlorophenol-indophenol (DCPIP) was added as electron acceptor to a final concentration of 70 µM. The reaction was started after 2 min pre-incubation by addition of succinate (20 mM, pH 7.4). Reduction of DCPIP was monitored at 600 nm ^37^. To account for background reactions, the assay was performed a second time in the presence of the SDH inhibitor malonate (20 mM) and the obtained activity subtracted from the first measurement.

Malate dehydrogenase (MDH) activity was determined following the oxidation of NADH to NAD^+^ at 340 nm. 20 µg ml^-1^ protein in Tris buffer (50 mM, pH 7.4) containing 100 mM NaCl were used. 250 µM NADH were added, and the reaction started by addition of oxaloacetate (2 mM) ^37^.

All enzyme activity assays were performed at 30°C and the initial slope during the linear change in absorption was used for calculating enzyme activity.

### Enzyme activity after plasma treatment

Bacteria were cultivated aerobically at 37°C in 15 ml LB or M9 medium as indicated until the cultures reached an OD_600_ of 0.6. Cells were harvested by centrifugation and resuspended in 400 µl LB medium. 20 µl of this highly dense bacterial suspension were applied onto a filter paper and exposed to the effluent of the µAPPJ for 60 s. For control samples, neither gas flow nor plasma were turned on. For gas control samples, cells were exposed to the helium/oxygen gas flow only. To accrue enough cell material, for each biological replicate of each strain, four filter papers were prepared, treated, and immersed together in one reaction tube filled with 0.9% (w/v) NaCl. The tubes were shaken vigorously for 2 min. The filter papers were then removed, and the samples were either used directly (t_0_) or incubated at 37°C for 20 min (t_20_) prior to further use to allow the cells to repair any treatment-induced damage. The samples (t_0_ and t_20_) were centrifuged, and the cell pellets resuspended in Tris buffer (50 mM, pH 7.4, 100 mM NaCl). Afterwards, the cells were lysed by sonication and extracts used to determine enzyme activities as described above.

### Determination of iron and cysteine concentrations

Intracellular iron was quantified using chromazurol S (CAS) ^38, 39^. *E. coli* wild type was incubated in M9 minimal medium supplemented with different amounts of FeSO_4_. Na_2_SO_4_ was added to yield a final sulfate concentration of 1 mM in all samples. At OD_600_ = 0.3, cells were harvested by centrifugation, washed thrice with NaCl (0.9% w/v) and finally resuspended in acetate buffer (200 mM, pH 4.7). Cetyltrimethylammonium bromide and CAS were added to final concentrations of 190 µM and 66 µM, respectively. After incubation at room temperature for 10 min, the absorption at 630 nm was determined. The standard curve for the calculation of intracellular ferric ion concentrations was generated using FeSO_4_ solutions of defined concentrations in the assay.

Intracellular cysteine concentrations were determined colorimetrically after derivatization of cysteine with ninhydrin ^40^. Strains were grown in 50 ml LB medium to an OD_600_ of 0.3. The cells were harvested, resuspended in 50 µl Tris buffer (50 mM, pH 7.4) and lysed by heating at 95°C for 10 min. Cell debris was removed by centrifugation. Dithiothreitol was added to a final concentration of 5 mM and samples were incubated at room temperature for 10 min for reduction of cystines. To 50 µl of the sample, 50 µl acetic acid (100%) and 50 µl of a freshly prepared ninhydrin solution (25 mg ninhydrin in 600 µl acetic acid (100%) and 400 µl HCl (37%)) were added. The samples were heated at 100°C for 10 min and directly afterwards cooled in an ice/water bath. The reaction product was conserved by addition of 100 µl ethanol (100%) until absorption at 560 nm was determined. Cysteine samples of defined concentrations were treated same way to generate a standard curve.

## 3. Results

### [FeS] clusters and plasma sensitivity

Biosynthesis and maintenance of [FeS] clusters is strictly regulated in bacterial cells. The two biosynthetic pathways existing in *E. coli*, the *isc* and the *suf* pathway, are interwoven on a regulatory level. To assess if [FeS] cluster homeostasis is important for survival of plasma treatment, single gene deletion strains impaired in different steps of [FeS] cluster synthesis and repair were exposed to the plasma effluent of a µAPPJ (**Fig. 1**). Survival of the wild type was set to 1.0. Several, but not all mutants coped less well with plasma treatment than the wild type. The Δ*sufC* strain showed the lowest relative survival rate (0.39) of all strains tested. Deletion of either *sufB* or *sufD* also resulted in increased plasma sensitivity, albeit to a lesser extent than deletion of *sufC*. All three proteins form the scaffold protein complex SufBC_2_D that is responsible for the assembly of [FeS] clusters from sulfur and iron under stress conditions ^41^. The equivalent enzyme to SufC in the *isc* pathway is IscU. The relative survival rate of the Δ*iscU* strain was similar to that of the wild type (0.98). Further differences between homologs of the two operons were observed for SufES and IscS. Both proteins are cysteine desulfurases and supply the sulfur for cluster assembly using cysteine as the source ^42^. The Δ*iscS* strain had a relative survival rate of 0.44, while the Δ*sufS* and Δ*sufE* strains showed rates of 0.79 and 1.1, respectively. Survival rates of Δ*sufA*, Δ*hscA*, Δ*fdx*, and Δ*iscX* strains under plasma exposure were similar to that of the wild type. The strains with an increased plasma sensitivity were analyzed regarding their growth in LB medium. Compared to the wild type, some of these mutants showed a growth defect in LB medium in the absence of any stress (**Supp.** Fig. 1). However, the growth defects did not correlate with differences in plasma sensitivity as determined by survival rates after plasma treatment (**Fig. 1**).

**Fig. 1.**
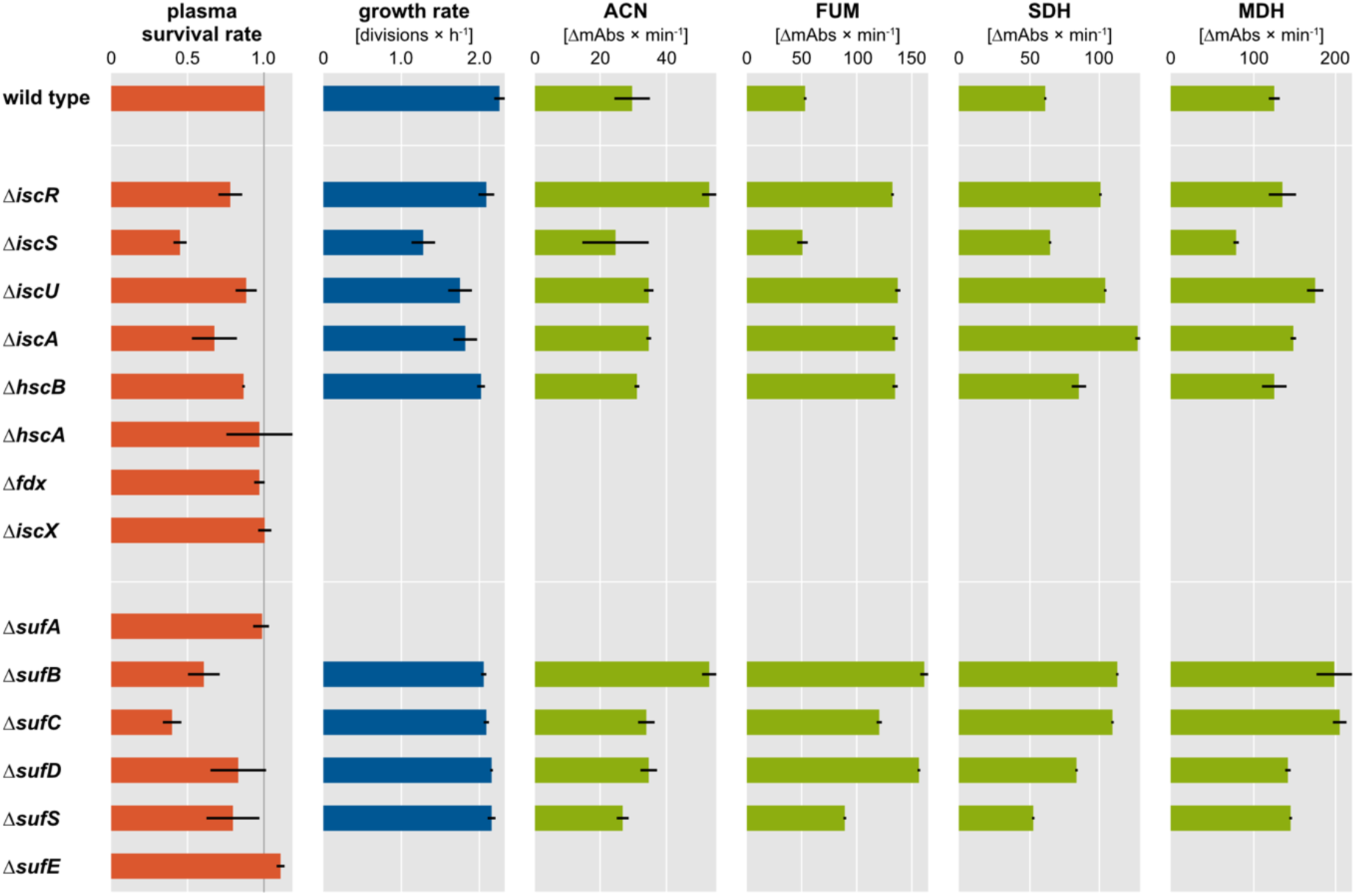
Characterization of strains with gene deletions in the *isc* or *suf* operon. Plasma survival rate: Deletion strains were exposed to the effluent of the µAPPJ for 30 s and colony forming units were determined. CFU of plasma-treated and gas-treated cells were set in relation to yield the survival rates. Strains with a plasma survival rate different from the wild type were further characterized. Growth rate: Single-gene deletion strains and the wild type were grown in LB medium and growth rates were determined during log-phase (between min 120 to 180 after inoculation) (see Fig. S1). Activity of [FeS] cluster proteins: Logarithmically growing deletion strains were lysed and enzyme assays were performed for aconitase (ACN), fumarase (FUM), succinate dehydrogenase (SDH), and malate dehydrogenase (not containing [FeS] clusters). Means and standard deviation of three independent biological replicates are shown.

The fact that several deletion strains exhibited lower plasma survival rates than the wild type (especially the Δ*sufB*, Δ*sufC*, Δ*iscA*, and Δ*iscS* strains) indicated an importance of [FeS] cluster homeostasis under plasma exposure. We hypothesized that the plasma-sensitive mutants either suffer globally from insufficient activity of [FeS] cluster-containing proteins making them hypersensitive to plasma treatment, or that [FeS] cluster-containing proteins are subject to rapid inactivation during plasma treatment and replacement and repair becomes critical under plasma treatment conditions. To test this, enzyme activity assays were established for [FeS] cluster-containing and [FeS] cluster-free enzymes. As surrogates for the cellular [FeS] cluster functionality, we chose three [FeS] cluster-containing enzymes from the citric acid cycle, aconitase (ACN), fumarase (FUM), and succinate dehydrogenase (SDH). Malate dehydrogenase (MDH), which is also part of the citric acid cycle, does not harbor any [FeS] clusters. The activities of ACN, FUM, and SDH were first used to assess the [FeS] cluster homeostasis in unstressed mutants (**Fig. 1**). To this end, the *isc* and *suf* deletion strains were grown to exponential phase, harvested, and enzyme activity assays performed on lysates. No direct correlation was observed when comparing the plasma sensitivity of the deletion strains to the activity of the [FeS] cluster proteins in unstressed mutants, indicating that the deletion strains are metabolically adapted to the specific genetic impairment in [FeS] cluster synthesis. *E. coli* has several proteins that catalyze the reactions monitored here as readout for the [FeS] cluster status. In *E. coli* there are two genes encoding aconitases, *acnB*, which is expressed under non-stress conditions, and *acnA*, which is less sensitive to oxidation and iron depletion ^43, 44^. There are also multiple fumarases, including FumC, an [FeS] cluster-free fumarase synthesized under O_2_^-^ stress ^45, 46^. The deletion strains likely adapted to their genetic impairment by adjusting expression levels and perhaps ratios between different complementary enzymes.

Next, we assessed the impact of plasma treatment on the activity of [FeS] cluster-containing proteins in the *E. coli* wild type. Cells were treated with only He/O_2_ gas flow or with the effluent of the µAPPJ for 1 min and then either harvested directly to perform activity assays or left to recover at 37°C for 20 min after plasma treatment to allow for restoration of [FeS] clusters before analyzing enzyme activity (**Fig. 2**).

**Fig. 2.**
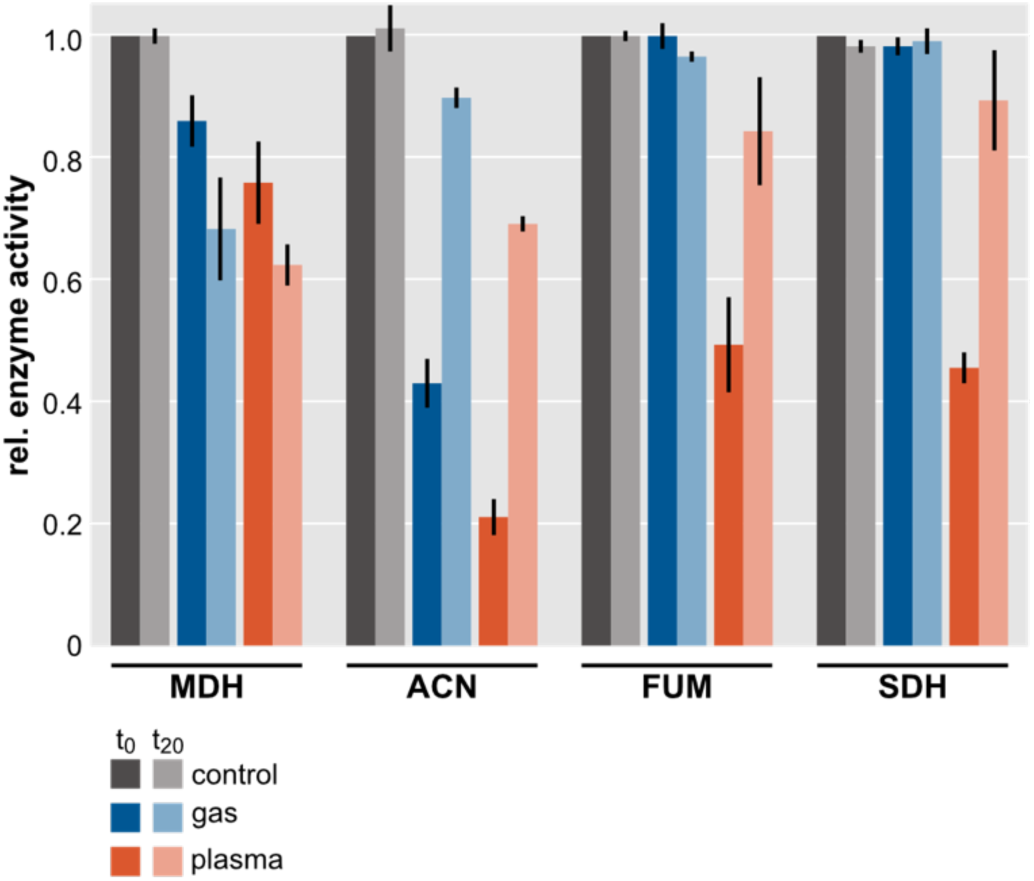
Enzyme activity of malate dehydrogenase (MDH), aconitase (ACN), fumarase (FUM), and succinate dehydrogenase (SDH) after µAPPJ treatment for 1 min. *E. coli* wild type cells were left untreated (gray, set to 1.0), exposed to gas flow (blue), or treated with plasma (orange). Residual enzyme activity was determined either directly after treatment (t0, dark bars) or after a 20 min post-plasma incubation at 37°C (t20, light bars). Averages and standard deviations represent three independent biological replicates.

MDH and ACN activities were reduced to 86% and 43%, respectively, in the gas-treated controls, which were exposed only to He/O_2_ mixture but not to plasma. When exposed to plasma, the activity of all three [FeS] cluster-containing enzymes was reduced approximately by half compared to gas treatment (that of aconitase was reduced by 79%), while MDH activity was only reduced by approximately 12%. The tested [FeS] cluster-containing enzymes are more susceptible to plasma-induced inactivation than enzymes without this cofactor. When cells were allowed to rest after plasma treatment at 37°C for 20 min (**Fig. 2**), enzyme activity was restored to a large degree. These data do not allow to differentiate between *de novo* synthesis, repair of [FeS] clusters, and upregulation of protein isoforms. Nevertheless, it is evident that plasma-treated cells are capable of overcoming plasma-induced damage to [FeS] cluster-containing enzymes or to bypass corrupted pathways.

### Restoration of [FeS] clusters

By allowing *E. coli* to regenerate after plasma treatment, activity of [FeS] cluster-containing enzymes was restored, albeit not to 100%. [FeS] clusters are synthesized from cysteine and iron under consumption of reducing agents (*e.g.* ferredoxin) ^41^. Changing cysteine and iron availability in the cells might change the regeneration capacities after plasma treatment and the overall plasma survival. Jaganaman *et al.* postulated that cysteine supplementation to LB medium is not sufficient to increase the sulfur availability for [FeS] cluster synthesis ^47^. It was shown that the reaction catalyzed by the serine acetyltransferase CysE is the bottleneck in cysteine biosynthesis ^48^. Thus, in this study, to increase the intracellular cysteine concentration, CysE was over-produced IPTG-dependently using the plasmid pCA24N::*cysE* ^32^. *cysE* expression was induced with different IPTG concentrations and growth rates, intracellular cysteine concentrations, plasma survival, and [FeS] cluster integrity monitored (**Fig. 3**, **Supp.** Fig. 2).

**Fig. 3.**
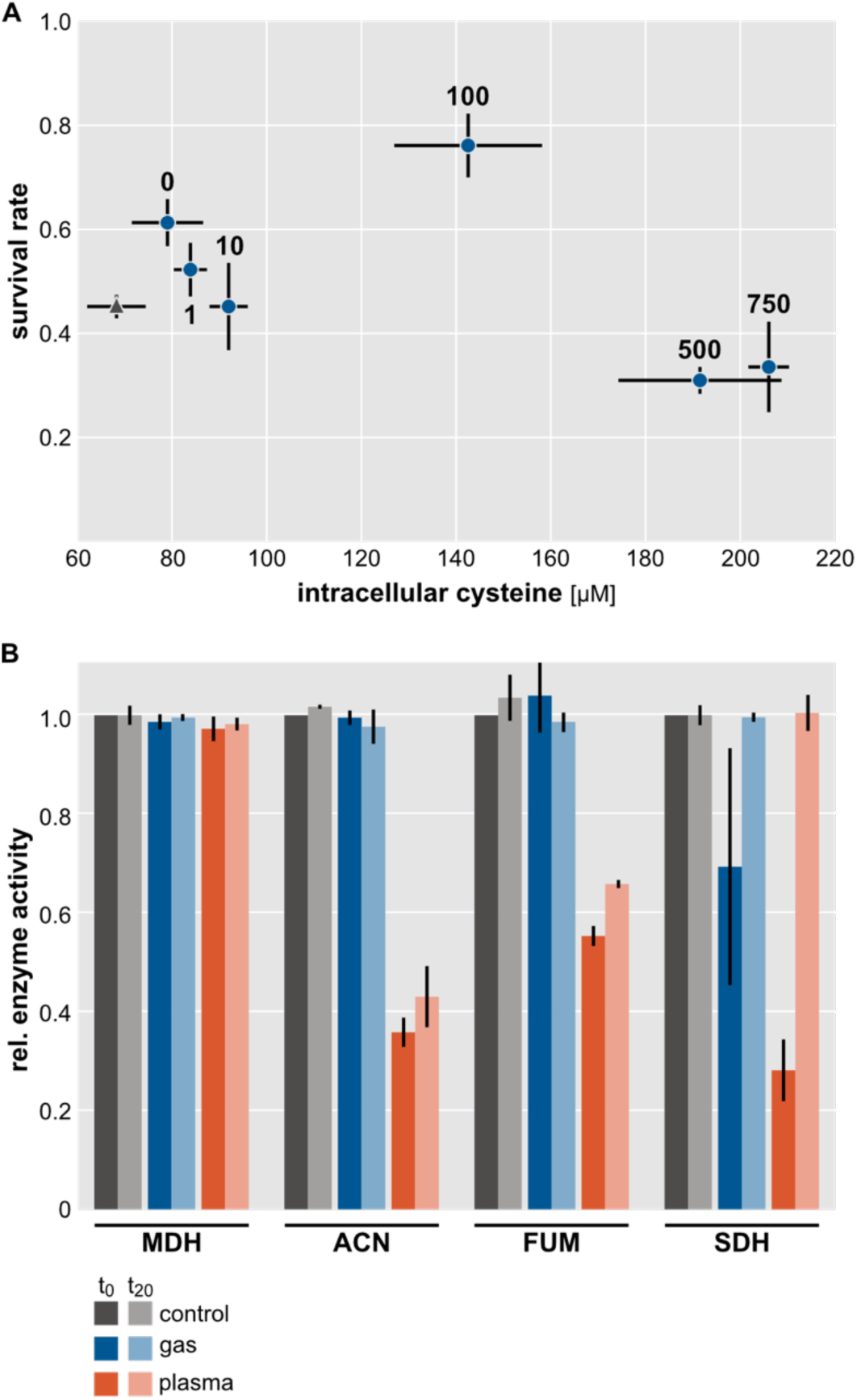
Influence of cysteine over-production. (**A**) Dependency of the plasma survival rate on the intracellular cysteine concentration in the strain *E. coli* pCA24N::*cysE* (blue). Amounts of IPTG for induction [µM] of *cysE* are indicated next to each data point. Data for *E. coli* wild type (without the plasmid) is given as gray triangle. (**B**) Enzyme activity of malate dehydrogenase (MDH), aconitase (ACN), fumarase (FUM), and succinate dehydrogenase (SDH) after µAPPJ treatment. *E. coli* pCA24N::*cysE* induced with 100 µM IPTG was left untreated (gray, set to 1.0), exposed to gas flow (blue), or treated with plasma (orange). Residual enzyme activity was determined either directly after treatment (t0, dark bars) or after incubation at 37°C for 20 min (t20, light bars). Averages and standard deviations represent three independent biological replicates.

In the absence of IPTG the intracellular cysteine concentration in the strain harboring the plasmid pCA24N::*cysE* was already increased to 79 µM compared to 68 µM in the plasmid-free wild type (**Fig. 3A**). This strain also had an increased survival rate of 0.6 (wild type: 0.44). With increasing IPTG concentrations cellular cysteine levels increased further, reaching 3-fold the cysteine concentration of the wild type at 750 µM IPTG. Higher cysteine concentrations did not generally correlate with increased survival rates. The culture induced with 100 µM IPTG and an intracellular cysteine concentration of 143 µM showed the highest plasma survival rate (0.75). The over-production of cysteine appears to present a burden for the cells as can be seen from the reduced growth rates of the strains at high IPTG concentrations (**Supp.** Fig. 2). High amounts of free intracellular cysteine have been shown to be cytotoxic by driving the Fenton reaction ^49^. Apparently, for plasma survival an optimal cysteine level exists around 140 µM.

In contrast to the wild type, in the *cysE* over-expressing strain the MDH and ACN activities were not affected by treatment with He/O_2_ gas (no plasma ignition). ACN, FUM, and SDH activities were affected by plasma treatment approximately to the same degree as in the wild type. And while the regeneration of ACN and FUM activities was slower than in the wild type, SDH activity was restored completely within the 20 min time window (**Fig. 3B**).

Next, iron availability was altered. To this end *E. coli* wild type was cultivated in M9 minimal medium supplemented with defined amounts of FeSO_4_ (**Fig. 4A**) ^47^. M9 minimal medium is the standard medium for growing *E. coli* according to the German Collection of Microorganisms and Cell Cultures (DSMZ) and the American Type Culture Collection (ATCC) and it contains no iron supplement. When adding defined amounts of FeSO_4_, the intracellular iron concentration increased 1.2-fold in cultures supplemented with 10 µM Fe^2+^. For this culture, the survival rate was 0.72 (without iron supplementation: 0.47). Higher amounts of intracellular iron did not further improve the survival rate after plasma treatment.

**Fig. 4.**
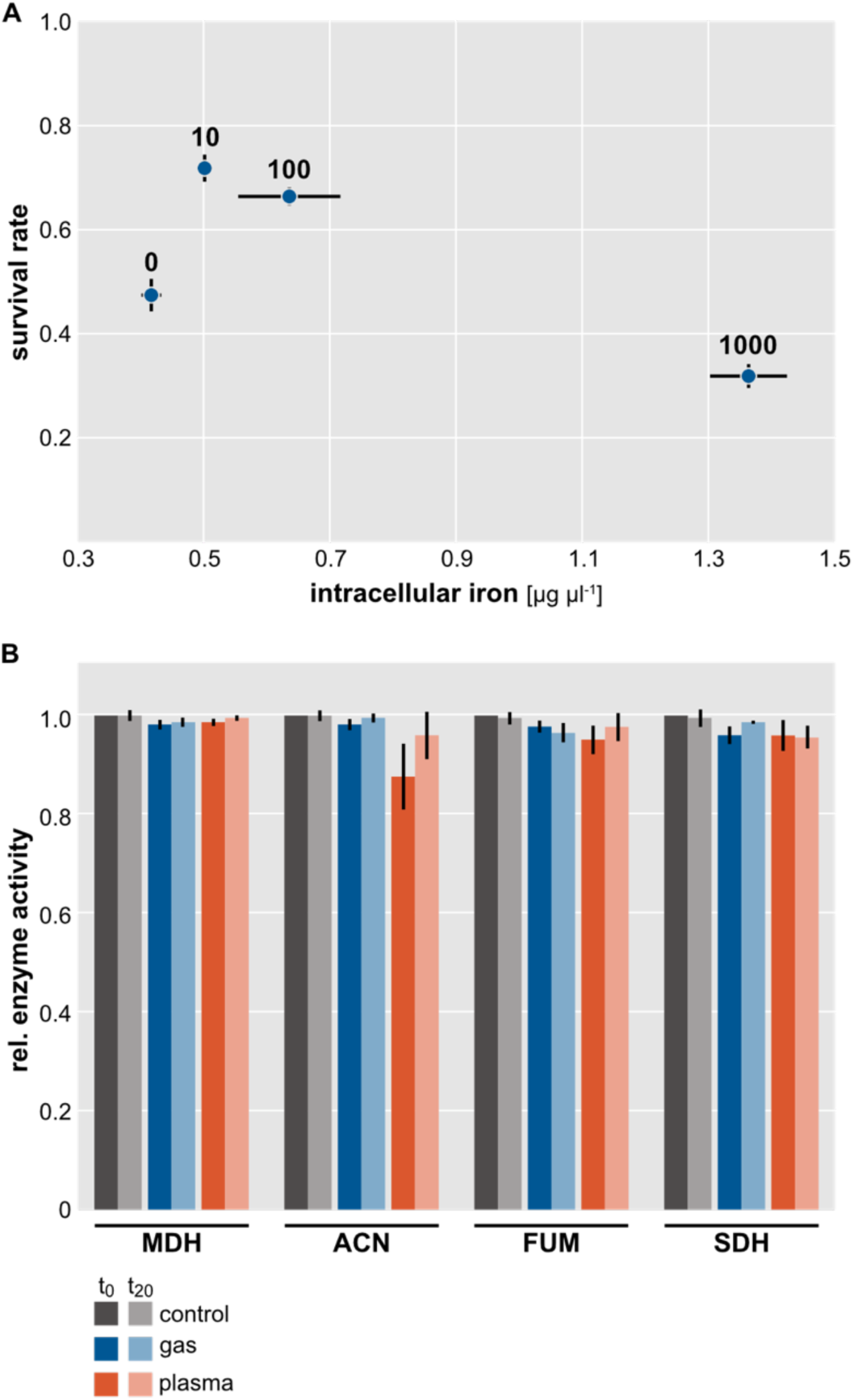
Influence of iron supplementation to the minimal medium. (**A**) Dependency of the plasma survival rate on the intracellular iron concentration in *E. coli* wild type. Amounts of FeSO4 [µM] added to the medium are indicated next to each data point. (**B**) Enzyme activity of malate dehydrogenase (MDH), aconitase (ACN), fumarase (FUM), and succinate dehydrogenase (SDH) after µAPPJ treatment. *E. coli* wild type supplemented with 10 µM Fe^2+^ were left untreated (gray, set to 1.0), exposed to gas flow (blue), or treated with plasma (orange). Residual enzyme activity was determined either directly after treatment (t0, dark bars) or after incubation at 37°C for 20 min (t20, light bars). Averages and standard deviations represent three independent biological replicates.

In M9 medium without iron supplementation, the [FeS] cluster-containing enzymes were protected from gas flow-induced damage, but as soon as plasma was ignited, their activity was reduced (**Supp.** Fig. 3B). In contrast to enhanced cysteine availability, in M9 medium without iron supplementation a rapid regeneration of ACN and FUM, but not SDH activity was observed. ACN and FUM both contain [4Fe-4S] clusters, while SDH harbors [2Fe-2S], [3Fe-4S], and [4Fe-4S] clusters ^27^. [FeS] clusters are damaged in different ways depending on the type of cluster, their role (Lewis acid or for electron transfer), and the protein surrounding, hence their repair presumably relies on different precursors. While regeneration of SDH activity seems to be limited by cysteine availability (**Fig. 3B**), ACN and FUM regeneration appears to be limited by a compound with higher availability in minimal medium.

When 10 µM FeSO_4_ was added to the minimal medium, enzyme activity was preserved not only when cells were exposed to He/O_2_ gas but also when cells were exposed to plasma (**Fig. 4B**), either due to a very efficient protection of the clusters or a very rapid repair.

### Superoxide as key species

[FeS] clusters are readily oxidized by superoxide causing their disassembly if not repaired ^29^. Moreover, superoxide was shown to be one of the most relevant plasma-generated species that bacteria need to be able to defend themselves from ^22^ and a deletion of superoxide-detoxifying enzymes caused cells to be more sensitive to plasma generated by a dielectric barrier discharge ^50^. We hypothesized, that superoxide dismutase activity is crucial for protecting [FeS] clusters in *E. coli* from plasma treatment. Single-gene deletion mutants of the three superoxide dismutases in *E. coli* (Δ*sodA*, Δ*sodB*, Δ*sodC*) as well as a double (Δ*sodAB*) and the triple deletion mutant (ΔSOD) were exposed to the effluent of the µAPPJ to determine their survival rates. (**Fig. 5A**). All three single deletion mutants exhibited a reduced survival rate (0.23-0.36, wild type: 0.54). Of the three, SodB seemed to be most important for bacterial survival under plasma treatment. SodB mainly protects cytosolic proteins from superoxide while SodA has been ascribed a role in DNA protection ^52^. In accordance with former studies, these data underscore the importance of plasma-induced protein damage as bacterial inactivation mechanism ^50^. The triple mutant (ΔSOD) was unable to survive plasma treatment. At least one superoxide dismutase was required for at least a small fraction of *E. coli* cells to survive.

**Fig. 5.**
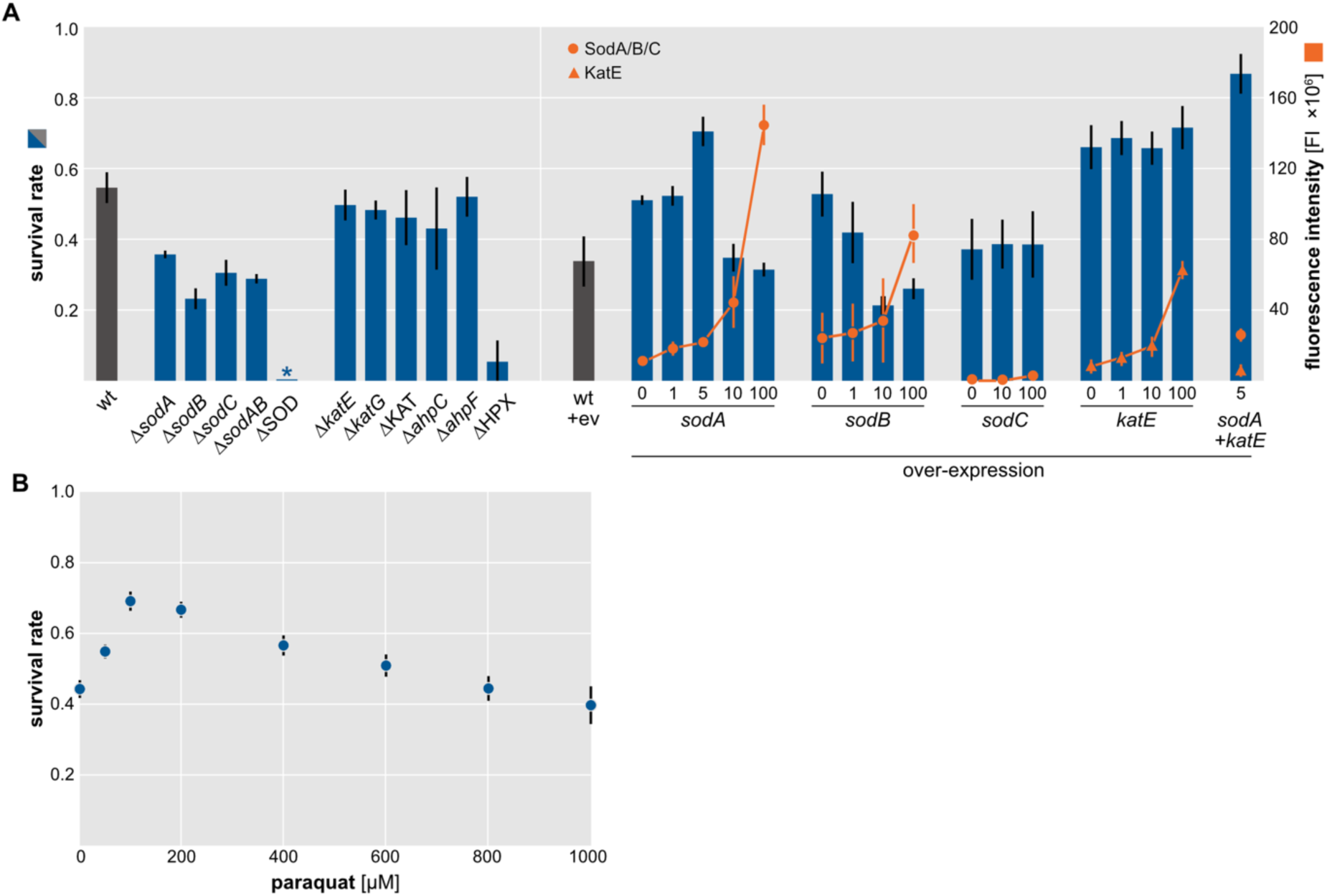
*E. coli* plasma survival. (**A**) Plasma resistance of gene-deletion or over-expression strains induced with IPTG [µM] at different concentrations. The asterisk indicates a survival rate of 0. Western blot analyses were conducted for quantification of the over-expression product shown by orange line graphs. An anti His6 antibody conjugated to a fluorophore was used for detection. (**B**) *E. coli* wild type was incubated with paraquat prior plasma treatment to allow for pre-adaptation. ev: empty vector pCA24N. Averages and standard deviations of three biological replicates are shown.

The three superoxide dismutases were over-produced individually, and plasma survival of over-production strains characterized (**Fig. 5A**). Over-production of SodA induced with 5 µM IPTG led to the highest survival rates (0.71, wild type harboring empty vector: 0.34). Higher concentrations of IPTG did not further improve survival despite higher SodA levels. For SodB, highest survival rates (0.53) were observed when no IPTG was added. No significant increase in plasma resistance was observed when inducing *sodC* over-expression by adding IPTG. Western blot analyses showed no increase in SodC production upon IPTG induction and it remains to be investigated if the N-terminal His-tag is being removed during translocation of the periplasmic SodC ^52^ or if no over-production of SodC takes place. Taken together, the experiments with deletion and over-expression strains indicate that a lack of SodB is most deleterious to the cells, while over-production of SodA is most beneficial. To test whether superoxide protection and plasma tolerance may be altered without genetic manipulation, simply due to physiological pre-adaptation, *E. coli* wild type was pre-incubated with paraquat, a redox-cycling agent, which generates superoxide (**Fig. 5B**) ^53^. Paraquat was added directly at inoculation of the main culture. Pre-incubation with 100 µM paraquat resulted in increased plasma tolerance (0.69). This experiment shows that the ability of bacteria to survive plasma treatment depends on the growth conditions.

### Protection of [FeS] clusters by superoxide dismutase and catalase

Over-production of SodA increased the survival rates after plasma treatment to 0.71, compared to 0.34 for the wild type harboring the empty vector. We investigated *in vivo* [FeS] cluster stability and regeneration after plasma treatment in the *sodA* over-expression strain induced with 5 µM IPTG (**Fig. 6**). While FUM activity was reduced to approximately 45% like in the wild type with empty vector, ACN and SDH activities were only slightly reduced indicating protection by SodA. It is known that superoxide is formed during plasma treatment ^53^. Plasma is a rich source of free electrons, which readily react with the oxygen admixed to the feed gas resulting in O_2_^-54, 55^. Furthermore, molecular oxygen is often the final acceptor of unpaired electrons from other radicals also yielding O_2_^-56^. The conversion of plasma-derived O_2_^-^ by SodA yields hydrogen peroxide, which is also a known toxicant of [FeS] clusters ^29, 57^. The cellular enzymes detoxifying hydrogen peroxide by converting it to water and molecular oxygen are the catalases. In *E. coli*, the proteins KatE, KatG, and AhpC_10_F_2_ build the first line of defense against H_2_O_2_. The deletion strains Δ*katE*, Δ*katG*, Δ*katE*Δ*katG* (ΔKAT), Δ*ahpC*, or Δ*ahpF* did not exhibited an increased plasma sensitivity, probably due to mutual complementation (**Fig. 5A**). However, the triple deletion mutant (Δ*katE*Δ*katG*Δ*ahpC*) ΔHPX had a very low plasma survival rate of 0.05.

**Fig. 6.**
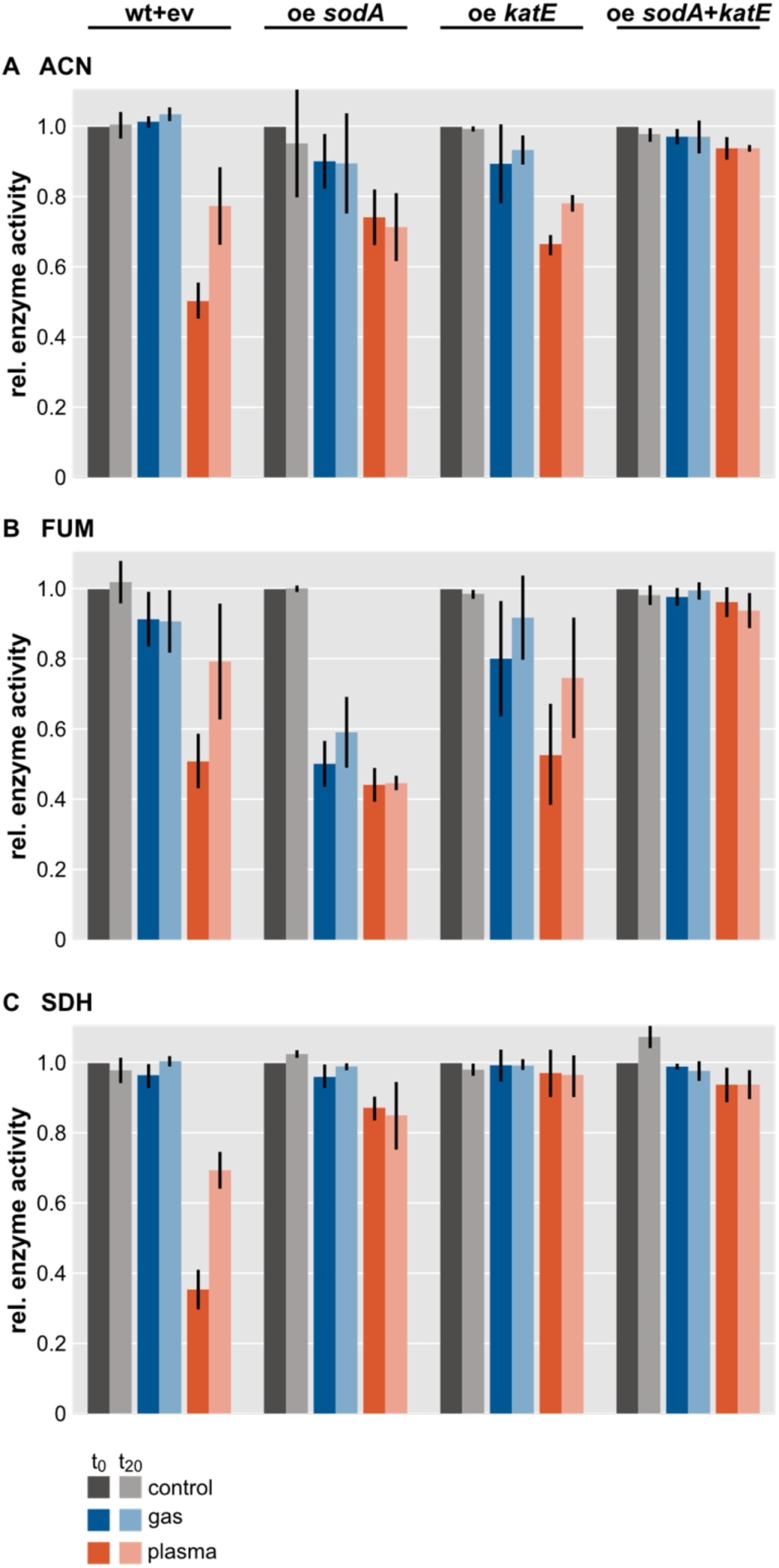
[FeS] cluster enzyme activity of *E. coli* over-expressing *sodA* and/or *katE*. Residual enzyme activity was determined either directly after 1 min plasma treatment (t0, dark bars) or after incubation at 37°C for 20 min (t20, light bars) in the following strains: *E. coli* wild type harboring the empty vector pCA24N (wt+ev), *E. coli* pCA24N::*sodA* (5 µM IPTG) (oe *sodA*), *E. coli* pCA24N::*katE* (5 µM IPTG) (oe *katE*), and *E. coli* pCA24N::*sodA*::*katE* (5 µM IPTG) (oe *sodA*+*katE*). Activity of aconitase (A, ACN), fumarase (B, FUM), and succinate dehydrogenase (C, SDH) was determined. The activity of the malate dehydrogenase is shown in Supp. Fig. S4. Averages and standard deviations represent three independent biological replicates.

In order to protect the [FeS] cluster enzymes from hydrogen peroxide accumulation in a *sodA* over-expressing strain, the gene *katE* was cloned into the plasmid pCA24N::*sodA* yielding the bi-cistronic expression plasmid pCA24N::*sodA*::*katE*. In the strain combining SodA and KatE over-production, all [FeS] cluster proteins were protected completely from the plasma-generated species by (**Fig. 6**). *katE* over-expression alone also improved the survival rate of *E. coli* (to 0.72) (**Fig. 5A**). Especially SDH activity was fully preserved by KatE over-production, and minor benefits were observed for ACN and FUM (**Fig. 6**). In SDH, the [FeS] clusters act as redox-cycling cofactors. They are likely inactivated by H_2_O_2_ as has been described for mitochondrial SDH ^58^. In ACN and FUM the clusters are Lewis acids with direct exposure to the solvent and, more sensitive to superoxide than to H_2_O_2_ ^59^.

To rule out that the over-production of protein led to unspecific scavenging of reactive species simply by the presence of high amounts of cytoplasmic protein acting as unspecific scavenger, Western Blot analyses were performed (**Fig. 5A**). The amounts of SodA were the same in the *sodA* and the *sodA*::*katE* expressing mutants, while the amount of KatE was notably lower in the *sodA*::*katE* expressing mutant compared to the *katE* expressing strain. This indicates that the combined catalytic capacities of the two detoxifying enzymes (and not the over-production of protein in general) protects the cells from the bactericidal effects of plasma.

## 4. Discussion

[FeS] clusters are the most abundant prosthetic group in proteins and participate in a wide range of reactions. Nevertheless, they evolved before oxygenation of the atmosphere and hence are prone to damage by various reactive oxygen species, such as O_2_^-^ and H_2_O_2_ ^29^. Based on a genome-wide analysis of plasma-sensitive *E. coli* mutants ^22^, considering their importance in many metabolic pathways, and their high sensitivity to known plasma-derived reactive species, we hypothesized that [FeS] cluster containing enzymes are good candidates for presenting the molecular weak spot when it comes to bacteria surviving plasma treatment.

### Inactivation of [FeS] cluster-containing enzymes

Different proteins have been exposed to atmospheric-pressure plasma treatment and typically a rapid inactivation was observed ^13, 18, 60–62^. In this study, [FeS] cluster-containing enzymes of a central metabolic pathway (citric acid cycle) were shown to be more sensitive to plasma-induced inactivation than an enzyme of the same pathway, that does not have an [FeS] cluster cofactor. Upon oxidative disruption of [FeS] clusters in a cellular context, more reactive oxygen species form and iron ions are released, which cause further damage by Fenton-like reactions ^63^. This chain reaction is known from oxidative killing of bacteria in phagolysosomes during the respiratory burst ^59, 64^. Since during plasma treatment some of the same reactive species are produced as by neutrophils and macrophages, plasma-mediated [FeS] cluster disruption may also initiate a bactericidal chain reaction.

Many reactive oxygen or nitrogen species are known to cause [FeS] cluster disassembly. A highly complex mixture of reactive species can be detected in plasma-treated liquids, so that the question arose as to which plasma species are responsible for [FeS] cluster enzyme inactivation in plasma-treated bacteria. The plasma jet was operated with a He/O_2_ mixture. The fact that [FeS] clusters of three enzymes could be completely protected by the joint over-expression of *sodA* and *katE* indicates that under these treatment conditions O_2_^-^ and H_2_O_2_ are the most relevant species for [FeS] cluster inactivation. Plasma-generated reactive nitrogen species such as ^•^NO play a very minor role, if any, in the plasma-induced inactivation of [FeS] clusters. The interaction of ^•^NO with [FeS] clusters has been described in literature ^65, 66^, but only low amounts of ^•^NO are detected in the effluent of the µAPPJ under the described operating conditions ^6^. Whether O_2_^-^ or H_2_O_2_ is more relevant for plasma-mediated [FeS] cluster inactivation cannot be inferred from the data generated in the presented experiments. The [FeS] clusters in SDH are responsible for the electron flow from succinate to ubiquinone ^27^ and are buried deep inside the protein, whereas the clusters in ACN and FUM are acting as Lewis acid exposing at least one iron ion to the solvent ^67^. SDH activity was fully protected in the *katE* over-expression strain (**Fig. 6**), indicating that the [FeS] clusters of SDH are inactivated by hydrogen peroxide. For full protection of ACN and FUM, the over-production of both SodA and KatE was required (**Fig. 6**). [FeS] clusters acting as Lewis acid seem to be vulnerable to both superoxide and hydrogen peroxide. Apparently for any [FeS] cluster, the ROS-mediated inactivation mechanism has to be elucidated individually ^58, 59^.

Importantly, the simultaneous over-production of SodA and KatE prevented cluster disruption completely and bacterial survival rates surpassed those of *sodA* or *katE* over-expressing strains. Whether or not [FeS] cluster-containing proteins present the weakest link in bacterial survival of plasma treatment or just a weak link, remains to be shown in future studies. Another candidate for the weakest link is glyceraldehyde 3-phosphate dehydrogenase (GAPDH), which exhibited a similar inactivation rate as the three [FeS] cluster-containing enzymes analyzed here ^17^. The *in vivo* GAPDH activity was reduced by more than 90% within a 60 s exposure to the effluent of the plasma jet used here. The activity of GAPDH depends on a catalytic cysteine residue known to be highly sensitive to oxidation, which was shown to be oxidized irreversibly by plasma treatment rendering the enzyme inactive ^17^.

### Repair of [FeS] clusters after plasma treatment

The activity of the [FeS] cluster-containing enzymes was partially restored within 20 min at 37°C. In the wild type, the residual enzyme activity doubled during regeneration (**Fig. 2**). ACN, FUM, and SDH employ [4Fe-4S]^2+^ clusters for catalysis. Those clusters are easily oxidized resulting in [3Fe-4S]^+^ clusters, which further decompose to [2Fe-2S]^2+^ clusters ^68^. The loss of one iron ion occurs most frequently ^69^. Depending on the kind and degree of damage, either iron or sulfur is needed for the repair of the clusters. In a cysteine over-producing strain, SDH activity was restored completely after plasma treatment (**Fig. 3B**), indicating that repair of the clusters relies on cysteine-derived sulfur.

There are limits as to the cysteine concentration that is advantageous for surviving plasma treatment (**Fig. 3A**), the reason for which is to be elucidated. High concentrations of cysteine are known to be cytotoxic as they promote the Fenton reaction ^49^. Thiol groups like those of cysteine are major targets for oxidation by plasma-generated species, leading to accumulation of oxidized and nitrosylated cysteine derivatives with unknown impact on cellular metabolism ^21, 70^. We considered that the effect might also be indirect and rooted in the fact that cells burdened with a very high over-production of a protein, such as CysE in this case, may be more sensitive to stress in general and plasma stress in particular. However, this does not seem to be the case generally, since not all protein over-expressing strains showed an increased plasma-sensitivity (see, e.g. the *sodA*, *katE* over-expressing strain). We conclude from our results that a moderate cysteine over-production increases the cellular capacity to repair plasma-damaged [FeS] clusters and increase survival rates.

A similar effect was observed for iron. Up to a point, elevation of the iron concentration in the growth medium resulted in increased survival rates and a complete protection of [FeS] clusters from plasma-mediated integrity loss (**Fig. 5**). Higher iron concentrations provided less of a benefit. Benov and Fridovich described a tolerance towards superoxide generated from paraquat, when *E. coli* cells were incubated in iron-rich medium. This benefit was attributed to scavenging of superoxide by [FeS] clusters, which protects the cells from further damage, and a prompt and efficient repair of oxidized clusters ^71^.

### Clinical relevance

Atmospheric-pressure plasmas are applied in clinical settings for instance in dermatology for treatment of chronic wounds. A potential unintended emergence of and selection for plasma-resistant bacteria is neglected ^72^, although a rapid development of resistance has been documented for antibiotic therapy and the application of biocides. Based on the present study, three possible mechanisms leading to temporarily or permanently plasma-insensitive bacteria can be charted: (i) environmental conditions, (ii) metabolic adaptation, and (iii) constitutive over-production of detoxifying enzymes. Environmental conditions such as the availability of certain nutrients like iron were shown to affect the plasma tolerance of *E. coli* (**Fig. 4A**). Depending on the micro-environment of the bacteria, the conditions may favor bacterial survival to a yet unknown extent.

Metabolic adaptation may also contribute to a temporary plasma tolerance. The activity of all three [FeS] cluster-containing enzymes was reduced by plasma treatment, but substantial regeneration occurred within 20 min (**Fig. 2**). The upregulation of repair machineries and of alternative pathways less sensitive to plasma-mediated inactivation could render a second treatment less effective. The *suf* operon, for example, is less susceptible to inactivation by oxidative stress than the *isc* operon ^73^. Such an adaptation could be triggered *e.g.* by plasma treatment of infected wounds, the pre-exposure to reactive oxygen and nitrogen species generated by the immune system, or – as shown here for the paraquat (**Fig. 5B**) – the exposure to reactive oxygen species in general. It can be expected that a faster repair and better protection of [FeS] clusters based on metabolic pre-adaptation will result in a transient increase in plasma tolerance.

Genetic mutations may permanently lead to increased plasma resistance. The over-expression of *sodA* alone led to a significant change in the survival rate of *E. coli* and the combined over-expression of *sodA* and *katE* increased the plasma resistance even further. Similar changes were already observed for other genes, albeit not to the same extent ^22^. Over-expression of a single gene can occur quite easily, *e.g.* due to mutations in the promoter region ^74^. It is worth keeping in mind that for many clinical antibacterial applications of plasma such as wound antisepsis, an intermediary plasma resistant would be detrimental, since human cells also suffer at long plasma exposure times ^75, 76^.

In sum, several mechanisms can lead to plasma insensitive bacteria, either permanently by genetic alterations or transiently by adaptation.

## 5. Conclusion

This study provides evidence that iron-sulfur clusters are among the molecular structures inside bacterial cells that are most prone to inactivation by atmospheric-pressure plasma, and therefore decisive for bacterial inactivation or survival. The stability of [FeS] cluster-containing enzymes was increased by altering iron availability in the growth medium, over-production of detoxifying enzymes (superoxide dismutase SodA and catalase KatE), or by pre-adaptation to reactive species. Temporary and permanent adaptation led to efficient protection of [FeS] cluster-containing enzymes, resulting in increased plasma survival rates.

## Acknowledgements

The authors thank Lars Leichert for the fruitful discussions and Jan Benedikt for technical advice. JEB gratefully acknowledges funding from the German Research Foundation (BA 4193/7-1, CRC1316, and RTG2341).

