## Supplementary Information for "Iron-sulfur cluster proteins present the weak spot in plasma-treated *Escherichia coli*"

#### Title

### Supplementary Figures

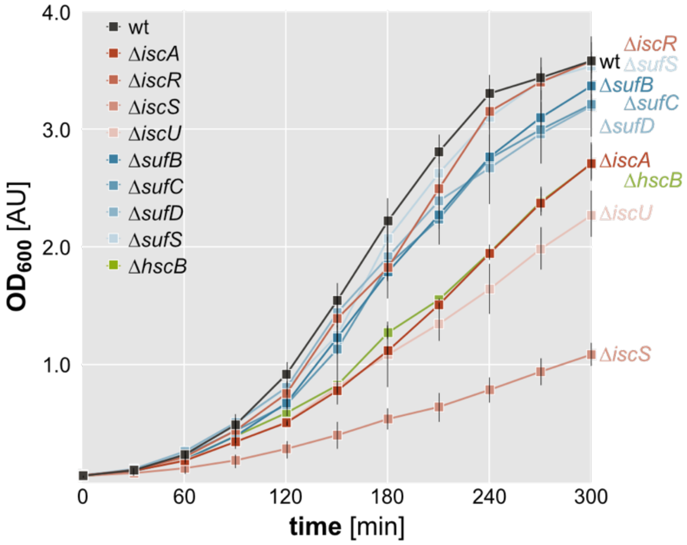


**Fig. S1.** Growth of *E. coli* wild type and deletion mutation in LB medium at 37°C. Averages and standard deviations represent three independent biological experiments.

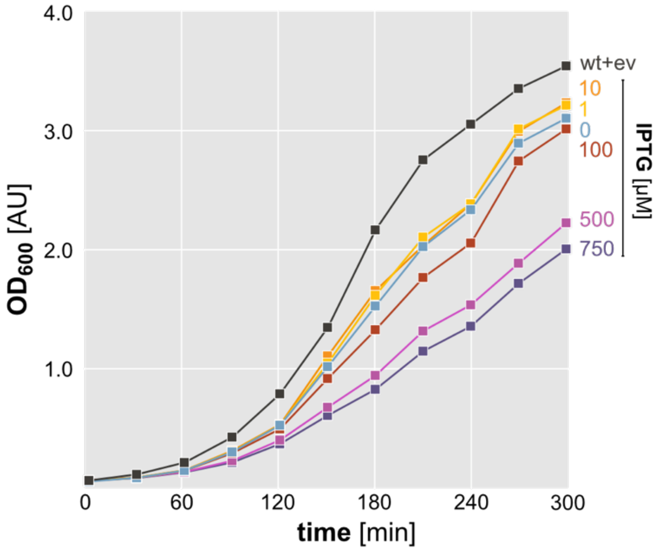


**Fig. S2.** Growth of *E. coli* wild type harboring the empty vector pCA24N (wt+ev) or a *cysE* over-expression plasmid (pCA24N::*cysE*) induced with different amounts of IPTG.

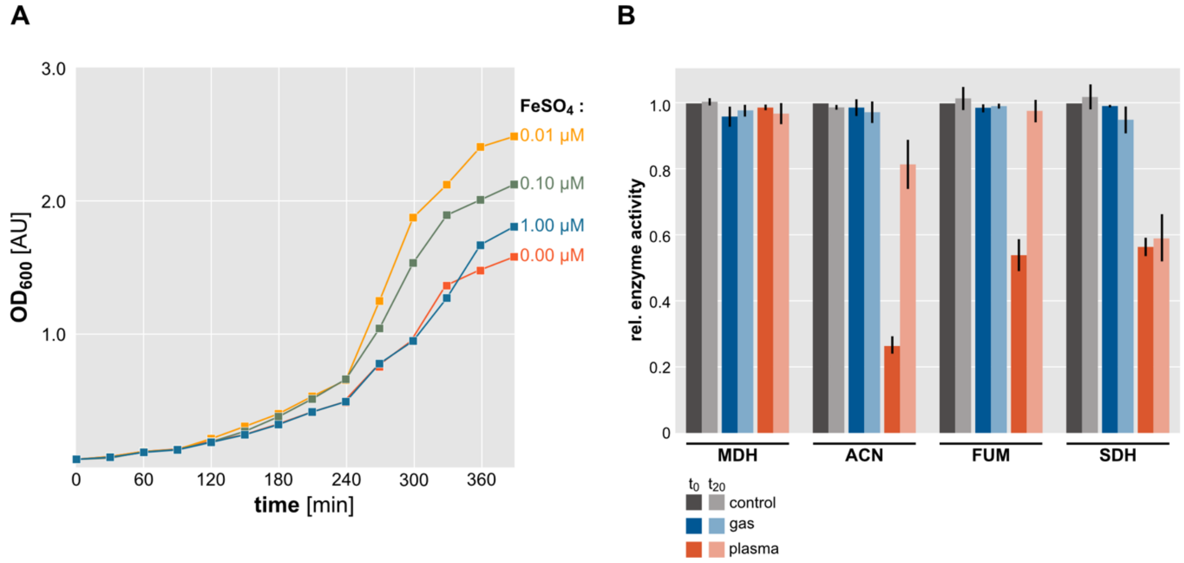


**Fig. S3.** (**A**) Growth of *E. coli* wild type in M9 minimal medium supplemented with $FeSO_4$. (**B**) Enzyme activity of malate dehydrogenase (MDH), aconitase (ACN), fumarase (FUM), and succinate dehydrogenase (SDH) after µAPPJ treatment. *E. coli* wild type grown in M9 medium without any iron supplementation was untreated (gray, set to 1.0), exposed to helium/oxygen gas flow (blue), or treated with plasma for 1 min (orange). Residual enzyme activity was determined either directly after treatment ($t_0$, dark bars) or after incubation at 37°C for 20 min ($t_{20}$, light bars). Averages and standard deviations represent three independent biological replicates.

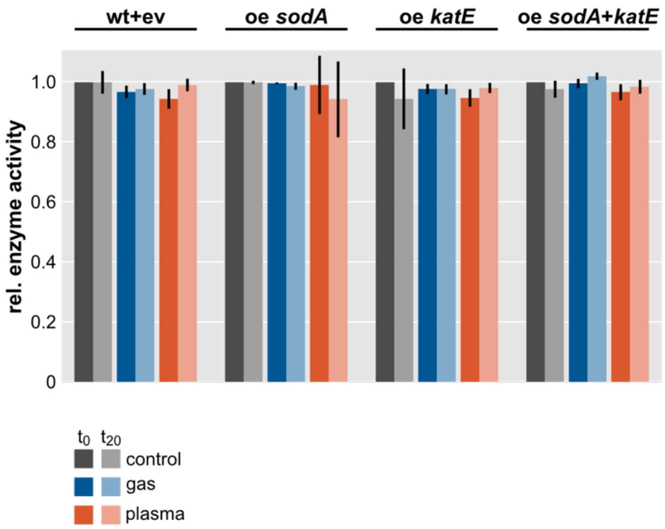


**Fig. S4.** Enzyme activity of malate dehydrogenase after µAPPJ treatment. Residual enzyme activity was determined either directly after 1 min plasma treatment ($t_0$, dark bars) or after incubation at 37°C for 20 min ($t_{20}$, light bars) in the following strains: *E. coli* wild type harboring the empty vector pCA24N (wt+ev), *E. coli* pCA24N::*sodA* (5 µM IPTG) (oe *sodA*), *E. coli* pCA24N::*katE* (5 µM IPTG) (oe *katE*), and *E. coli* pCA24N::*sodA*::*katE* (5 µM IPTG) (oe *sodA+katE*). Averages and standard deviations represent three independent biological replicates.
